# Temporal Dynamics of Brain Activity Predicting Sense of Agency over Muscle Movements

**DOI:** 10.1101/2023.05.06.539706

**Authors:** John P. Veillette, Pedro Lopes, Howard C. Nusbaum

**Affiliations:** Department of Psychology, University of Chicago 60637; Department of Computer Science, University of Chicago 60637

**Author notes:** Correspondence should be addressed to John P. Veillette.

## Abstract

Our muscles are the primary means through which we affect the external world, and the sense of agency (SoA) over the action through those muscles is fundamental to our self-awareness. However, SoA research to date has focused almost exclusively on agency over action outcomes rather than over the musculature itself, as it was believed that SoA over the musculature could not be manipulated directly. Drawing on methods from human-computer interaction and adaptive experimentation, we use human-in-the-loop Bayesian optimization to tune the timing of electrical muscle stimulation so as to robustly elicit a sense of agency over electrically-actuated muscle movements in male and female human subjects. We use time-resolved decoding of subjects’ EEG to estimate the time course of neural activity which predicts reported agency on a trial-by-trial basis. Like paradigms which assess SoA over action consequences, we found that the late (post-conscious) neural activity predicts SoA. Unlike typical paradigms, however, we also find patterns of early (sensorimotor) activity with distinct temporal dynamics predicts agency over muscle movements, suggesting that the “neural correlates of agency” may depend on the level of abstraction (i.e., direct sensorimotor feedback vs. downstream consequences) most relevant to a given agency judgement. Moreover, fractal analysis of the EEG suggests that SoA-contingent dynamics of neural activity may modulate the sensitivity of the motor system to external input.

**Significance Statement:** The sense of agency – the feeling of “I did that” – when directing one’s own musculature is a core feature of human experience. We show that we can robustly manipulate the sense of agency over electrically actuated muscle movements, and we investigate the time course of neural activity that predicts the sense of agency over these actuated movements. We find evidence of two distinct neural processes – a transient sequence of patterns that begins in the early sensorineural response to muscle stimulation and a later, sustained signature of agency. These results shed light on the neural mechanisms by which we experience our movements as volitional.

## 1. Introduction

Voluntary movements are usually accompanied by an experience of “I did that.” This feeling is the sense of agency (SoA), which is considered a basic building block of conscious selfhood (Gallagher, 2000; Haggard, 2008). Pathologies affecting SoA – including schizophrenia (Frith, 2012), alien hand syndrome (Panikkath et al., 2014), perceived (non-)control of a phantom limb (Ramachandran and Hirstein, 1998), automatic “utilization behavior” (Lhermitte et al., 1986), and learned paralysis (Wolf et al., 1989) – are often characterized by anomalies in the experience of control over the body itself (i.e., the musculature) rather than external action outcomes per se.

SoA research in healthy populations, however, has focused primarily on external consequences of action (Haggard, 2008, 2017). While some studies have manipulated bodily agency by delaying visual feedback from movements, such manipulations leave intact the somatic sensation of muscle movement, over which the subject might still feel agency in the absence of SoA over the decoupled visual stimulus (Tsakiris et al., 2010; Abdulkarim et al., 2023). Others have noted the lack of experimental paradigms addressing “narrow” SoA over muscles as opposed to “broad” SoA encompassing action outcomes (Christensen and Grünbaum, 2018). The field often assumes conclusions drawn from paradigms investigating SoA over a tone following a button press will generalize to other classes of agency judgments. As such, the literature tends to treat SoA as a homogenous phenomenon always accompanied by the same neural correlates. However, an alternative hypothesis is that the neural correlates of SoA may vary as a *function of modality* (e.g., proprioceptive vs. auditory) or the *level of abstraction* for a given judgement (Charalampaki et al., 2022). Indeed, it has been argued current models may not generalize to SoA over the musculature (Christensen and Grünbaum, 2018) or over thoughts (Frith, 2012). This discrepancy bears on a fundamental question of whether mechanisms that give rise to the experience of an agentic self are common across scales of biological and social-behavioral organization – and if not, how and why do we assign agency to the same unified self at these different scales (Veillette et al., 2023a)?

One reason for the shortage of paradigms assessing SoA over movements is that control over one’s own muscles is normally unambiguous. Indeed, previous attempts to elicit SoA for experimenter-evoked (e.g., by TMS) movements have not succeeded (Haggard and Clark, 2003; Christensen and Grünbaum, 2018), leading to the conclusion that “involuntary movements are never accompanied by a sense of agency” (Haggard, 2017). However, since cognitive scientists favor indirect SoA measures such as intentional binding (perceived delay) between actions and outcomes, these findings primarily reflect SoA over outcomes rather than SoA over the muscles (Haggard et al., 2002). Meanwhile, human-computer interaction researchers have begun investigating SoA in interfaces that use electrical-muscle stimulation (EMS) to drive users’ muscles. They find, in contrast, that subjects report EMS-caused movements as self-caused so long as stimulation temporally aligns with users’ endogenous intention to move (Kasahara et al., 2019, 2021; Tajima et al., 2022). Cognitive neuroscientists have yet to embrace these findings, partly because self-reports may result from response biases (Dewey and Knoblich, 2014), lacking convergent validation from neural measurements.

Thus, in the present work, we “preempt” subjects’ endogenous movements with EMS during a cue-response reaction time task, using manipulating stimulation timing to control the proportion of EMS-caused movements perceived as self-caused. Using time-resolved decoding of subjects’ trial-by-trial EEG, we show that cortical activity predicts agency judgements about resulting muscle movements as early as 83 ms following stimulation, showing that subjects’ self-report has a basis in early, pre-conscious sensorimotor processing – not just a response bias. Notably, this result differs from those obtained using typical button-tone paradigms, where early evoked responses (to tones) have repeatedly failed to predict subjective agency judgments (Kühn et al., 2011; Timm et al., 2016). Finally, an exploratory analysis shows that fractal measures also predict SoA, suggesting that complexity of sensory processing may differ for sensations perceived as movement feedback.

## 2. Materials and Methods

### 2.1. Methods Summary

The goal of our experimental design was to evoke movements using electrical muscle stimulation (EMS) in which roughly 50% of such movements were perceived as self-caused (agency) and 50% as EMS-caused (non-agency), even though all such movements are in fact EMS-caused. Previous work has shown that EMS-caused movements are perceived as self-caused if they modestly preempt subjects’ natural movements in a reaction time task (Kasahara et al., 2019; Tajima et al., 2022), and varying the timing of stimulation has been used to manipulate agency (Kasahara et al., 2021). However, this manipulation results in the stimulation latencies being systematically different between agency and non-agency trials, presenting a clear confound for neural analysis. To this end, we designed a procedure in which stimulation timing is tuned on a per-subject basis to a latency at which subjects report (without further manipulation of stimulation latency) that movements were self-caused on approximately half of trials (see *2.3* below). This results in maximally similar distributions of stimulation latency across agency and non-agency trials.

In our analysis, we aim to identify patterns in the scalp EEG response to muscle stimulation that robustly predict whether resulting muscle movements are perceived as self-caused or perceived as EMS-caused on a trial-by-trial basis, with a particular focus on the temporal characteristics of those patterns. Firstly, we train a linear classifier at each time (relative to stimulation onset) throughout the epoch, and test its generalization performance across subjects and across time (see *2.4.2*). An advantage of this approach is that it gives us information not just about when patterns that predict agency emerge, but how long those pattern remains present and continually predictive (King and Dehaene, 2014). This allows us to differentiate, for instance, patterns of neural activity that appear only transiently from those that are sustained over time. In addition, we assess whether complexity measures of the EEG response – the fractal dimension, and index of signal complexity, and the Hurst exponent, and index of long-range temporal dependency – predict trial-by-trial SoA. These complexity features allow us to uncover some of the qualitative characteristics of neural dynamics (e.g. sensitivity to perturbation, scale-freeness or self-similarity) in the presence and absence of SoA, though it should be noted since this latter analysis was exploratory, its evidential value should be weighted accordingly.

### 2.2. Participants and Ethics Statement

25 University of Chicago undergraduate students (6 male, 19 female, ages 19-24) participated in the study; however, two subjects were subsequently excluded for noncompliance with task instructions (i.e., one admitted to letting the electrical stimulator perform the task without attempting a volitional button press, and another pressed the button continually to speed through the task instead of when cued). Participants were recruited through the University of Chicago’s human subject recruitment system, SONA Systems. All subjects gave written, informed consent before participating. All of the methods performed in the study were in accordance with relevant safety and ethics guidelines for human subject research and were approved by the Social and Behavioral Sciences Institutional Review Board at the University of Chicago (IRB19-1302). This study was not a clinical trial.

### 2.3. Experimental Design

Subjects completed three blocks of trials: a pretest block (30 trials), a stimulation block (250 trials), and a posttest block (30 trials). After initiating each trial, subjects waited for a visual indicator to cue their movements (see Figure 1b). After the visual indicator was triggered (2-4 seconds, uniformly distributed after trial start), subjects attempted to press a button (on a *Cedrus RB-620* button box; California, United States) as quickly as possible. After the button was pressed, the trial ended.

**Figure 1:**
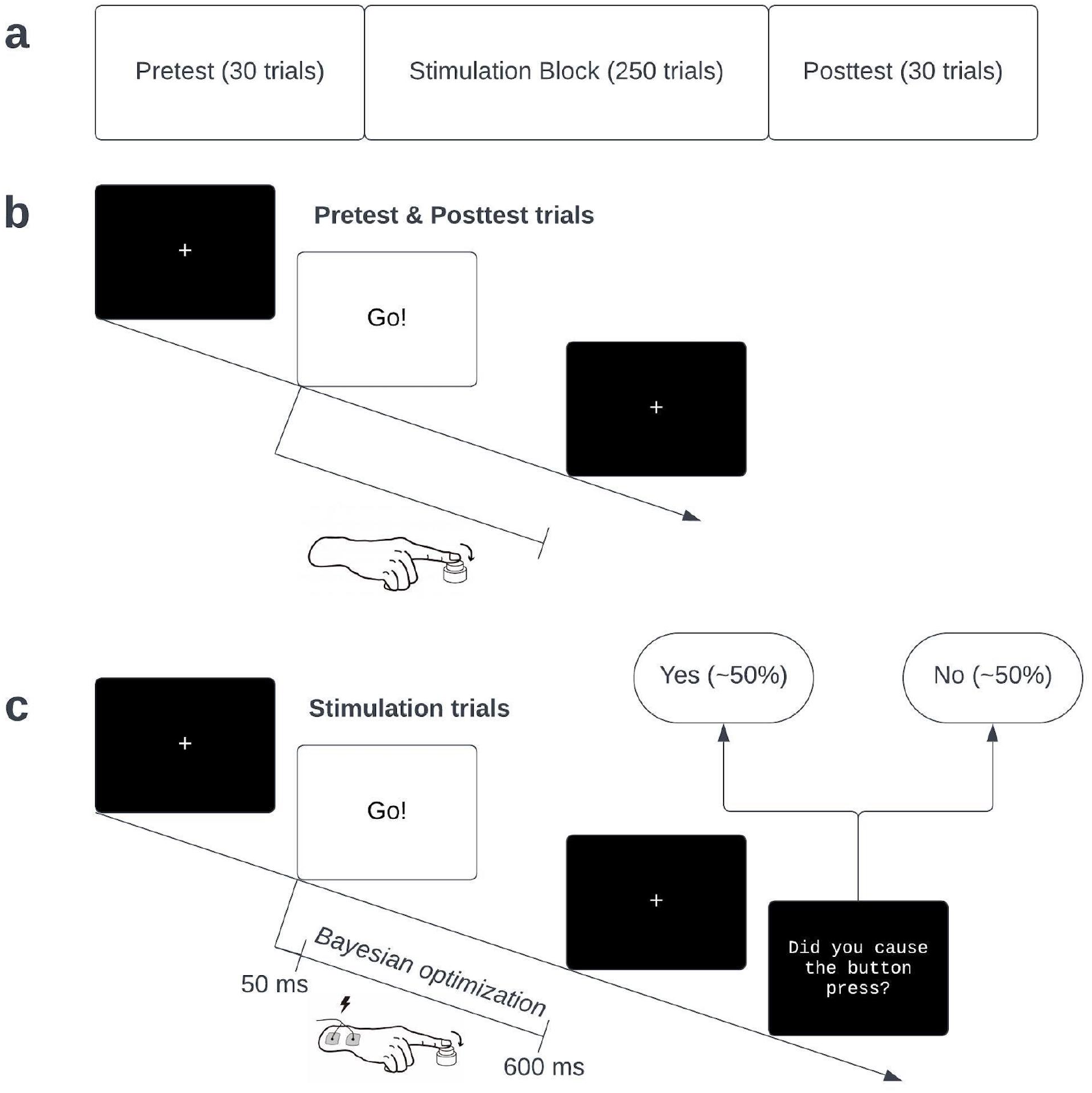
Task design. (a) Full experiment consists of a short pretest to gauge subjects’ reaction times, a stimulation block, and a posttest block to ensure true reaction times did not change dramatically over the course of the experiment. (b) Trials follow a typical cue-response reaction time paradigm, in which subjects are asked to press a button as quickly as possible following a cue to move. (b) In the stimulation block, subjects still attempt the reaction time task, but their natural movements can be preempted by muscle stimulation. After each trial, subjects guess whether the muscle movement resulting in the button press was self-caused or caused by muscle stimulation. Responses are used to tune the timing of muscle stimulation to a latency between 50-600 ms at which roughly 50% of trials are perceived as self-caused via Bayesian optimization (as shown in Figure 2).

During the stimulation block, however, electrical muscle stimulation (EMS) was applied to the forearm after the cue to move, with the aim of preempting subjects’ self-caused movement with an EMS-caused movement (see Figure 1c). After each trial, subjects were asked to report whether they caused the movement (agency) or the EMS caused the movement (non-agency).

Stimulation timing was adjusted on a trial-by-trial basis using a Bayesian optimization procedure designed to apply EMS as close as possible to the stimulation latency at which subjects would report agency with 50% probability (see Figures 1c, 2). Specifically, after each trial, we fit a Bayesian logistic regression predicting the probability of SoA from stimulation latency with a log-normal prior on the 50% threshold, centered 40 ms before subject’s mean reaction time observed during the pretest block and a log-normal (and thus constrained to be positive) slope prior. This 40 ms prior on preemptive timing was based on that reported in previous work (Kasahara et al., 2019, 2021). Each trial’s stimulation latency was drawn from the posterior distribution (truncated between 50-600 ms post-cue) of the 50% threshold in the logistic function fit after the previous trial. If the subject pressed the button before stimulation was delivered, stimulation occurred immediately upon the button press so that the onset of the electrical stimulation, which causes a perceptible though painless tingling sensation on the skin, was always temporally confusable with that of the movement. However, due to the speed at which the optimization procedure converges to reliably preempt subjects’ movements, such instances were quite rare (see *3.1* in *Results*).

**Figure 2:**
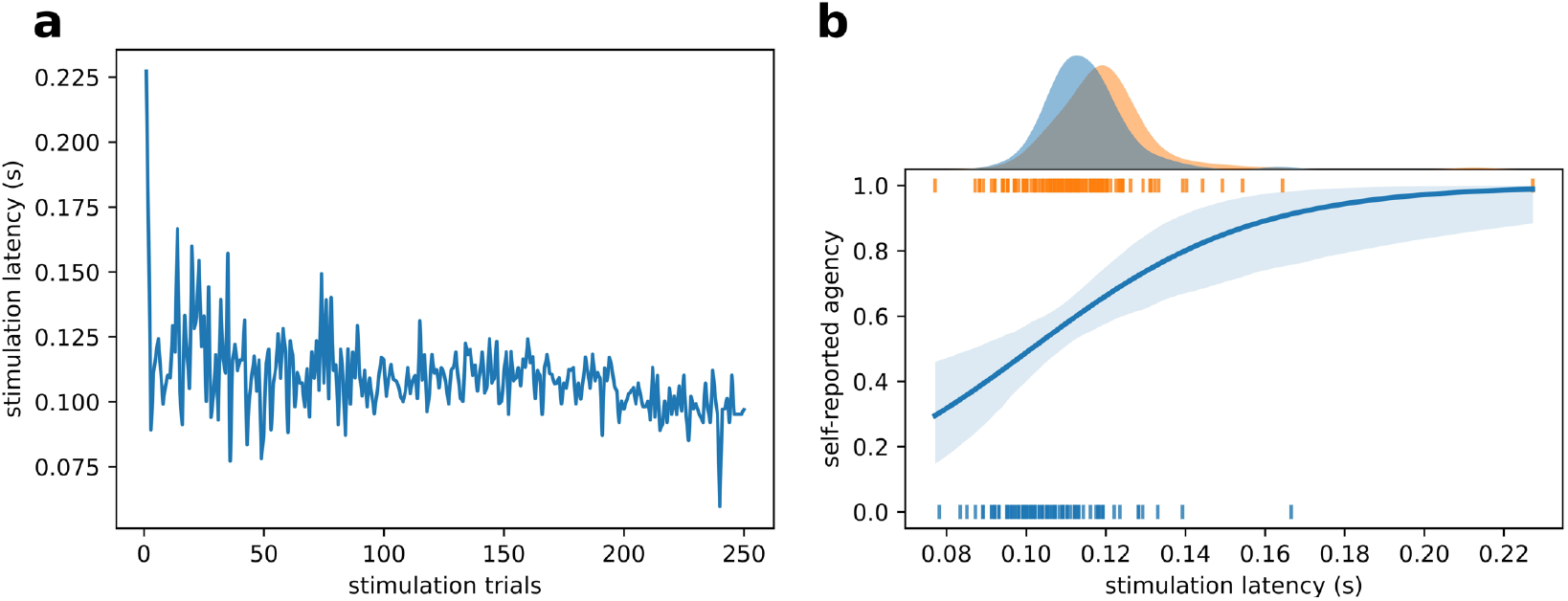
Trial-by-trial stimulation latency over the course of the stimulation block for a representative subject. (a) Stimulation latency hones in on a stable value over time, as a result of the Bayesian Optimization procedure. (b) A logistic regression computed after the experiment and shows that stimulation times are close to the retroactively estimated 50% threshold, even though that threshold was not known in advance. The subject featured here is “sub-07” in the associated dataset.

Visual stimulus presentation was implemented using PsychoPy (Peirce, 2007) and Bayesian optimization using the Python-embedded probabilistic programming language Pyro (Bingham et al., 2019). All code has been made available (see *2.7*).

### 2.4. Statistical Analysis

#### 2.4.1. Manipulation Checks and Outlier Removal

First, outlier removal was applied to remove trials in which muscle movement was not caused by electrical stimulation. Thus, we removed trials in which (a) subjects pressed the button prior to stimulation, (b) the recorded response time was outside the stimulation time window (i.e. greater than 600 ms), or (c) the lag between the EMS pulse and the corresponding button press fell outside of the middle 95% of the best fit log-normal distribution, indicating ineffective stimulation or the subjects’ endogenous movement coinciding with the EMS-caused movement. These steps removed an average of 30.8 trials per subject, after measured “reaction time” (button press) is a roughly linear function of the stimulation latency (see Figure 3b).

**Figure 3:**
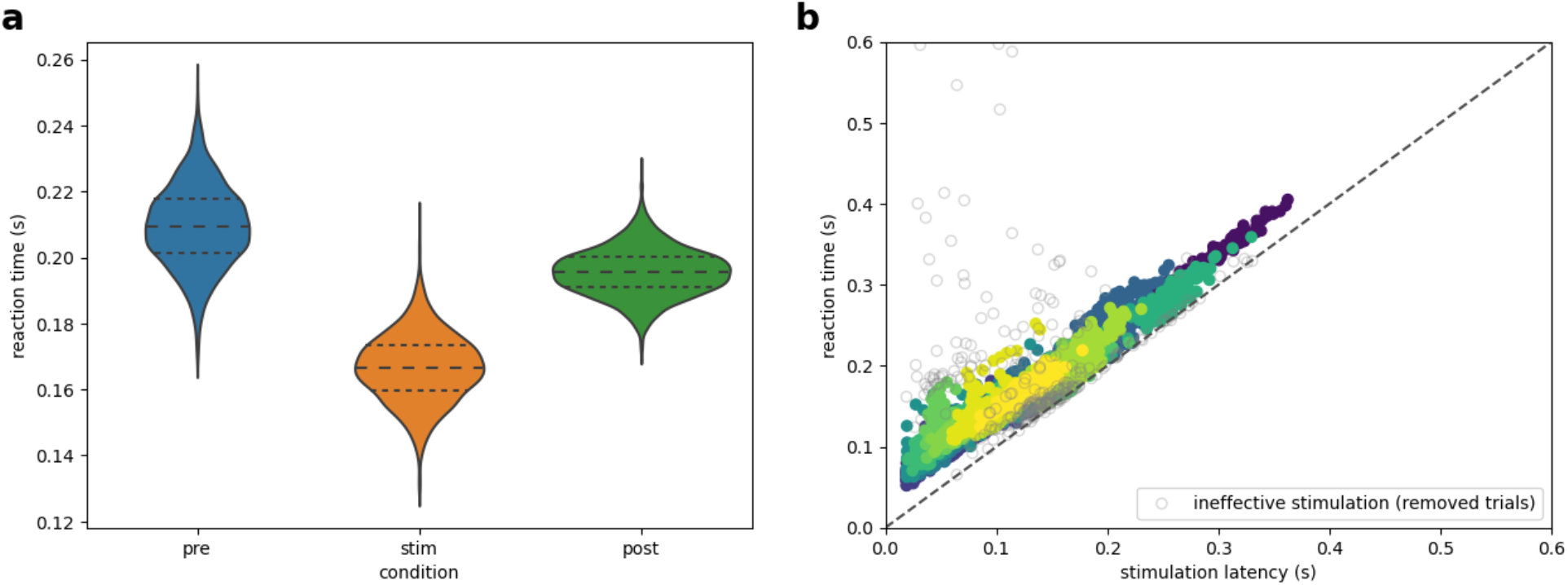
Electrical muscle stimulation consistently preempted subjects’ volitional movements. (a) Posterior distributions of the mean reaction times in each condition show that EMS-induced muscle movements occur earlier than subjects’ natural muscle movements. (b) After outlier removal, measured “reaction” times (shown for all trials and subjects) are a nearly linear function of the stimulation latency, indicating that movements in the remaining trials are, in fact, EMS-actuated.

To assess whether we were truly preempting subjects’ movements, we then fit a Bayesian multilevel model to the trial-by-trial reaction times in each experiment block with the *Bambi* package (Capretto et al., 2022) using the conservative default priors (Westfall, 2017). If we were preempting subjects’ movements, then reaction times in the stimulation block should be faster than in both the pretest or the posttest block.

As a final manipulation check, we then assessed whether stimulation latencies differed systematically between agency and non-agency trials. While prior work has shown that agency judgements vary as a function of stimulation latency (Kasahara et al., 2019, 2021; Tajima et al., 2022), the aim of our Bayesian optimization procedure was to minimize this confound by maximizing the overlap in stimulation latencies across agency and non-agency trials. Consequently, we fit a logistic regression predicting agency judgements from stimulation latency (with a random effect per subject, as in our EEG decoding analysis) to test whether any residual relationship between the two is strong enough to drive our EEG results.

#### 2.4.2. Linear EEG Decoding

After preprocessing of the EEG signal (described in *2.6*), we assess the temporal dynamics of patterns which differentiate agency and non-agency trials using the temporal generalization method (King and Dehaene, 2014). In this approach, a linear classifier is trained on the pattern of voltages at each timepoint (using a training set), and then its classification performance is quantified at every other timepoint (using a test set), yielding information about both the occurrence and duration of neural patterns which predict the outcome of interest.

In our case, we use a logistic regression (fit using generalized estimating equations to account for subject-level random effects) as a linear classifier (Liang and Zeger, 1986; Seabold and Perktold, 2010) to predict agency judgements, and we quantify classification performance using the area under the receiver-operator curve (ROC-AUC), a non-parametric, criterion-free metric of class separation. By “criterion-free” we mean that, unlike metrics such as accuracy which depend on a particular decision boundary, ROC-AUC reflects the tradeoff between false positives and false negatives across all possible decision boundaries; due to its weak assumptions, approximately normal distribution under the null hypotheses, and robustness to class imbalances, ROC-AUC is often recommended for multivariate pattern analyses of the EEG (King and Dehaene, 2014). Classification performance is calculated only on hold-out subjects (i.e. subjects not seen during classifier training), in a stratified 10-fold cross-validation scheme repeated 10 times. Cross-validated ROC-AUC scores are compared to chance performance (ROC-AUC = 0.5) using a one-sample *t*-test with a variance correction to account for non-independence between ROC-AUC values computed across cross-validation splits (Nadeau and Bengio, 1999).

This results in a 𝑛_times_ × 𝑛_times_ matrix 𝑀, where 𝑀_*ij*_ is the performance of the classifier trained at time 𝑖 evaluated at time 𝑗, as well as a *p*-value for each (*i, j*) pair. The “shape” of above-chance decoding performances can then be interpreted as providing information as to the temporal characteristic of predictive patterns of neural activity. For instance, if a pattern is predictive only on the diagonal (*i = j*), that pattern is transient. On the other hand, if classification performance remains above chance off-diagonal (*j > i*), then one can conclude the same pattern persists (and continues to predict SoA) across time. However, such conclusions are only licensed if one corrects for multiple comparisons using a method that allows inference about the “shape” of an effect, which common cluster-based corrections in the EEG literature do not (Sassenhagen and Draschkow, 2019). We use All-Resolutions Inference (Rosenblatt et al., 2018), which can compute simultaneous lower bounds on the true positive proportion in each cluster across all 𝑛_times_ × 𝑛_times_ possible clustering thresholds. This approach conveys uncertainty about the localization of true effects within clusters. For instance, if the proportion of true positives in a cluster is low, then one can conclude there is are true positive effects within the cluster but it is unclear precisely where; conversely, if the proportion is high (e.g., >95%), then the localization is quite certain.

#### 2.4.3. Complexity-based EEG Decoding

In this analysis, we assessed whether certain complexity measures of EEG response to stimulation – the fractal dimension and the Hurst exponent – could predict agency judgements. These metrics measure nonlinear properties of timeseries which can be used to inform qualitative claims about those timeseries’ underlying dynamics.

The fractal dimension, which we estimate using Higuchi’s algorithm (Higuchi, 1988), is a measure of the complexity or “roughness” of a time series (or of its underlying dynamical attractor). The fractal dimension of both the background EEG and the EEG response to perturbation is highly predictive of states of consciousness (Kesić and Spasić, 2016; Ruiz de Miras et al., 2019), consistent with some accounts of conscious awareness (Oizumi et al., 2014). Some preliminary evidence suggests that fractal dimension is higher for conscious percepts that are internally generated (e.g. mind wandering), making it a reasonable candidate predictor for sense of agency (Ibáñez-Molina and Iglesias-Parro, 2014). However, the interpretation of the fractal dimension on its own it ambiguous; it can be interpreted as reflecting how “self-similar” or “scale-free” a time series is, or alternatively as reflecting the local complexity of its dynamics. These interpretations can be disambiguated in the context of the Hurst exponent.

The Hurst exponent, which we estimate using rescaled range analysis, is a measure of long-range temporal dependencies in a time series (Qian and Rasheed, 2004). In the cognitive neuroscience literature, these long range dependencies have been argued to reflect how much local events – such as an external input – can alter the course of a neural system, assuming that events which substantially impact the system should have consequences which persist in time (Churchill et al., 2016; Kardan et al., 2020; Zhuang et al., 2022). Hurst is notably suppressed in those suffering from psychiatric conditions associated with an impaired sense of agency (Sokunbi et al., 2014; Stier et al., 2021).

If a time series is strictly self-similar, then the fractal dimension *D* will be related to the Hurst exponent *H* by the deterministic relationship 𝐷 + 𝐻 = 2 (Gneiting and Schlather, 2004), but these metrics have been reported to diverge in EEG data despite the 1/*f* power spectrum of resting/background EEG implying some degree of self-similarity in the signal (Martis et al., 2015). Unfortunately, estimates of the Hurst exponent computed from time-series as small as our single-trials are known to be biased (Oliver and Ballester, 1998; Eke et al., 2000, 2002). While the bias of our Hurst estimates prevents us from testing the 𝐷 + 𝐻 = 2 relationship directly, *within*-subject variation in trial-by-trial Hurst exponents estimated from EEG have been shown to be sensitive to cognitive functions (Kardan et al., 2020). Thus, if the EEG time series is self-similar or scale-free, then should the fractal dimension increase with agency, Hurst should decrease and vice versa. However, if they both positively or both negatively covary with agency (or one covaries and not the other), then the EEG response to stimulation is unlikely to be self-similar.

Since we perform this analysis at the electrode-level (instead of training a single classifier on all electrodes), we apply a current source density transformation before computing per-electrode complexity measures to increase the interpretability of spatial information in the EEG signal (Kayser and Tenke, 2015). For each electrode and subject, then, we compute the ROC-AUC between both of these metrics and subjects’ self-reported agency. Since there are no fit-to-the-data parameters in this analysis, no cross-validation scheme is necessary; a one-sample, two-sided *t-*test is used compare subject-level decoding performance to chance (ROC-AUC = 0.5).

### 2.5. Electrical Muscle Stimulation

Before the experiment began, two EMS electrodes were applied to the skin above the *flexor digitorum profundus* muscle on the right (dominant) forearm, which is an easily accessible muscle that moves the ring finger, which subjects used to press the button during the experiment. Stimulation was performed with a *RehaStim 1* device (*HASOMED* GmbH, Mageburg, Germany). On each trial, muscle actuation consisted of a single, biphasic pulse of constant current stimulation lasting 900 microseconds (400 μs positive, 400 μs negative, separated by 100 μs).

Before beginning the experiment, we calibrated the stimulation amplitude to the minimum intensity required to reliably move the subject’s finger. The calibration procedure was as follows: (a) The subject placed their ring finger on the button that would be used during the experiment and was instructed not to move their hand. (b) Starting at an intensity of 1 mA, we stimulated the subjects’ arm 10 times. If fewer than 10 button presses were registered, then we iterated the intensity by 1 mA and repeated. (c) We stopped increasing the intensity upon achieving 10 consecutive actuated button presses, or if a conservative safety limit of 25 mA was reached.

### 2.6. EEG Acquisition and Preprocessing

EEG was recorded with 64 active Ag/AgCl electrodes (*actiCHamp*, Brain Products, Munich, Germany) placed according to the extended 10-20 system. At the time of recording, the electrodes were referenced to Cz and sampled at 10,000 Hz. Two of the 64 electrodes (which would have been AF7 and AF8 on the typical *actiCAP* layout) were dropped below the left and right eyes so that they could later be re-referenced to become EOG channels. Experiment events were marked in the EEG recording using TTL triggers and later corrected with a photo-sensor (*Brain Products*, Munich, Germany) on the subjects’ screen. The precise subject-specific positions of the 62 head electrodes were measured at the end of each recording using a *CapTrak* (*Brain Products*, Munich, Germany).

EEG was later preprocessed in *Python* using *MNE-Python* package (Gramfort et al., 2014). First, we fit a multi-taper estimation of the sinusoidal components at the line noise frequency and its first two harmonics to partially attenuate electrical interference before interpolating the stimulus artifact. Then, the electrical artifact from the EMS pulse was removed by linearly interpolating over the interval starting 5 ms before and ending 10 ms after the event timestamp. Then, after interpolation, we applied an additional FIR notch filter at 60 Hz and its harmonics up to the intended upper passband edge (see below) to thoroughly clean the data of line noise, and then resampled the data to 5,000 Hz to improve the speed of computation for subsequent preprocessing steps.

Next, we applied common preprocessing operations in adherence with the standardized *PREP* preprocessing pipeline for EEG data (Bigdely-Shamlo et al., 2015) using the implementation in the *PyPREP* package (Appelhoff et al., 2022). This pipeline robustly re-references the EEG signal to the average of all electrodes and interpolates electrodes it determines have poor signal quality; see the PREP paper for a full description (Bigdely-Shamlo et al., 2015). A record of which channels were interpolated is available in subject-specific preprocessing reports (see ***2.7***).

Then, we filtered the data to the frequency band used for analysis. We used a single low cutoff of 1 Hz to remove low-frequency drift, but we used different high cutoff values for the different analysis described in *2.4.2* and *2.4.3* above. For linear decoding (*2.4.2*) we used a 30 Hz high cutoff; this filter setting is common for the analysis of event related potentials, as this level of temporal smoothing helps to align short neural events across subjects (Luck, 2014), and we posited such smoothing would likely improve sensitivity for between-subject decoding as we employ here. However, since fractal dimension is fundamentally a measure of signal roughness, which would be distorted by anything that would artificially smooth the signal, we used a more liberal 70 Hz high cutoff for the fractal analysis described in *2.4.3*.

We then removed EOG contamination of the EEG signal. We decomposed to EEG data into 15 independent components (ICs) using the *FastICA* algorithm (Hyvarinen, 1999). Then, we correlated each IC with the EOG channels, *z-*scored the correlation coefficients, and deemed an IC to contain eye artifact if the absolute value of its *z-*score exceeded 1.96. Those ICs were zeroed out to remove them from the original data. Plots of the scalp topographies of removed ICs for each subject can be found in their preprocessing reports (see ***2.7***).

Subsequently, we segmented the data into epochs starting 100 ms before the onset of stimulation and ending 500 ms after stimulation. We then estimated the peak-to-peak rejection threshold that would be used to identify trials containing unwanted artifacts using the *Autoreject* package (Jas et al., 2017), which estimates the optimal threshold as that which minimizes the 5-fold cross-validated root-mean-squared difference between the mean of the training folds and the median of the testing fold, a robust proxy metric for signal-to-noise. The resulting per-subject rejection thresholds are recorded in each subjects’ preprocessing report (see ***2.7***).

Since the visual evoked response to the movement cue is unlikely to be over by the time of stimulation, we attempted to remove the visual evoked response from our epoched data to minimize confounds. To do so, we computed evoked responses to both the visual and electrical stimuli simultaneously using a regression-based overlap correction on the continuous (non-epoched) data, excluding second-long chunks of the data in which peak-to-peak amplitude exceeds the rejection threshold (Smith and Kutas, 2015); conceptually, this is very similar to the way generalized linear models (GLMs) are used to deconvolve hemodynamic responses in fMRI. Then, the overlap-corrected visual evoked response was aligned with the epoched version of the data and subtracted out. Thus, the average visual response to the movement cue was removed from the stimulation-locked epochs. Subject-level evoked responses can be found in our open dataset and are visualized in the subject-specific preprocessing reports (see ***2.7***).

Finally, the rejection threshold was applied to the cleaned and overlap-corrected epochs, removing trials still contaminated by artifacts. The surviving epochs were down-sampled to twice their high-cutoff frequency for computational expediency and saved for further analysis. This epoched data is available in our open dataset, and subject-level trial yields are recorded in the accompanying quality check reports (see ***2.7***).

### 2.7. Data and Code Availability

Code for running the experiment can be found on *GitHub* (<u>github.com/apex-lab/agency-experiment</u>) and in a permanent archive on *Zenodo* (<u>doi.org/10.5281/zenodo.7894011</u>). Similarly, all data analysis code, including EEG preprocessing code, can be found at github.com/apex-lab/agency-analysis and https://doi.org/10.5281/zenodo.7894007. All data, including both raw data, preprocessed derivatives, and post-preprocessing quality check reports for each subject, can be found on *OpenNeuro* (<u>doi.org/10.18112/openneuro.ds004561.v1.0.0</u>).

### 2.8. Statistical Power

There is no widely agreed-upon approach for estimating the statistical power for detecting novel EEG effects, in which the spatiotemporal distribution of the effect is unknown a priori, as we recently reviewed (Veillette et al., 2023b). Statistical power for EEG effects depends not just on the number of subjects but also on the number of trials, and how these two design considerations interact to affect power seems to differ between components of the EEG response (Boudewyn et al., 2018; Hall et al., 2023; Jensen and MacDonald, 2023). However, statistical power for well-known EEG effects has been studied using a recently introduced Monte Carlo simulation approach (Boudewyn et al., 2018), and it is worth considering how well our study is powered for detecting effects reported in the literature. While we and others have found, using such a simulation-based approach, that a relatively small number of subjects and trials achieves very high statistical power for detecting the presence of seven endogenous EEG evoked response effects (Jensen and MacDonald, 2023; Veillette et al., 2023b), our main study result – that which differs from previous EEG studies of SoA – concerns an early (<200 ms) effect, and such effects usually reflect amplitude changes in exogenous response components present in both conditions rather than the presence or absence of an endogenous component. This more realistic case has been studied for three early evoked response components (Hall et al., 2023). Closest to our sample size, Hall and colleagues report that a within-subject design with a sample of 25 subjects, each having 120 trials per condition, achieves a power of at least 0.8 for detecting a 1.4 μV amplitude difference in the N1 component (in the window of 84-124 ms), a 1.3 μV difference in the Tb component (124-164 ms), and a 1.7 μV difference in the P2 component (151-191 ms) with a significance level of 0.05. Based on this comparison, we would expect our linear classifiers to be sensitive (i.e. with power of roughly 0.8) to amplitude differences on the order of ∼1.5 μV.

## 3. Results

### 3.1. Bayesian optimization effectively controls the proportion of trials perceived as self-caused

The Bayesian Optimization procedure resulted in trial-by-trial stimulation latencies honing in on some threshold estimate throughout the stimulation block. A representative time course is shown in Figure 2.

After removing trials in which stimulation failed to produce a muscle movement (and therefore the “reaction” time was not a function of stimulation latency), our multilevel model of the recorded reaction times estimated a 99.9% posterior probability that button presses occurred earlier in the stimulation block than in either of the other blocks. In particular, we estimate that “reaction” times resulting from EMS-actuated movements were between 17.5 ms and 65.0 ms faster than true reaction times in the first (pre) block with 95% probability, and between 13.8 ms and 43.9 ms faster than those in the final (post) block. A nominal speedup between the pre and post blocks was observed with 90.6% probability (95% HDI: [-6.7 ms, 32.8 ms]), suggesting that subjects may have improved their reaction times by the end of the task, but not enough to account for the much lower reaction times in the stimulation block. Posterior distributions for the (group) mean response times in each condition are shown in Figure 3. Taken together with the near linear relationship between stimulation latency and reaction time, we can conclude that movements were usually caused by muscle stimulation rather than the subject, effectively preempting subjects’ volitional movements.

While it is evident that muscle movements in the stimulation block (after outlier removal) were overwhelmingly caused by EMS rather than by the subject, subjects still reported that they caused roughly half of the movements. Overall, after outlier removal (see Figure 3), 51.98% of all trials across all subjects were judged as self-caused. On average, subjects reported that they caused 50.99% (SD: 14%) of movements. In other words, the Bayesian Optimization procedure was effective at controlling the proportion of trials in which movements were experienced as self-caused, generating a roughly 50-50 split of agency vs. non-agency trials.

While it is understood that agency judgements in this task paradigm vary as a function of the stimulation latency (Kasahara et al., 2019, 2021; Tajima et al., 2022), our Bayesian optimization procedure converges to a narrow latency range around the 50% agency threshold quickly enough to attenuate this confound. A logistic regression predicting agency judgements from stimulation latency (with a subject-level random effect) – notably the same approach we use to predict agency judgements from the EEG signal – fails to find a statistically significant relationship between the two (beta = 0.95, 95% CI: [-0.91, 2.82], p = 0.315). Thus, any residual relationship between stimulation latency and SoA is unlikely to explain our EEG findings (see below).

### 3.2. Distinct early and late neural processes predict agency judgments

Our linear decoding procedure showed above-chance decoding performance *across subjects*, reaching up to ROC-AUC = 0.587; thus, the patterns which we report predict agency judgements generalize across individuals. While we report the true-positive proportion within clusters across all clustering thresholds (see Figure 5b), we will focus primarily on the clusters in which the true positive proportion exceeds 95%, since these clusters are where we are sufficiently certain about the localization of the effect (Rosenblatt et al., 2018). The grand-average EEG evoked response to muscle stimulation is provided, for visualization only, in Figure 4; this may be useful context when considering predictive topographies, as shown in Figure 6.

**Figure 4:**
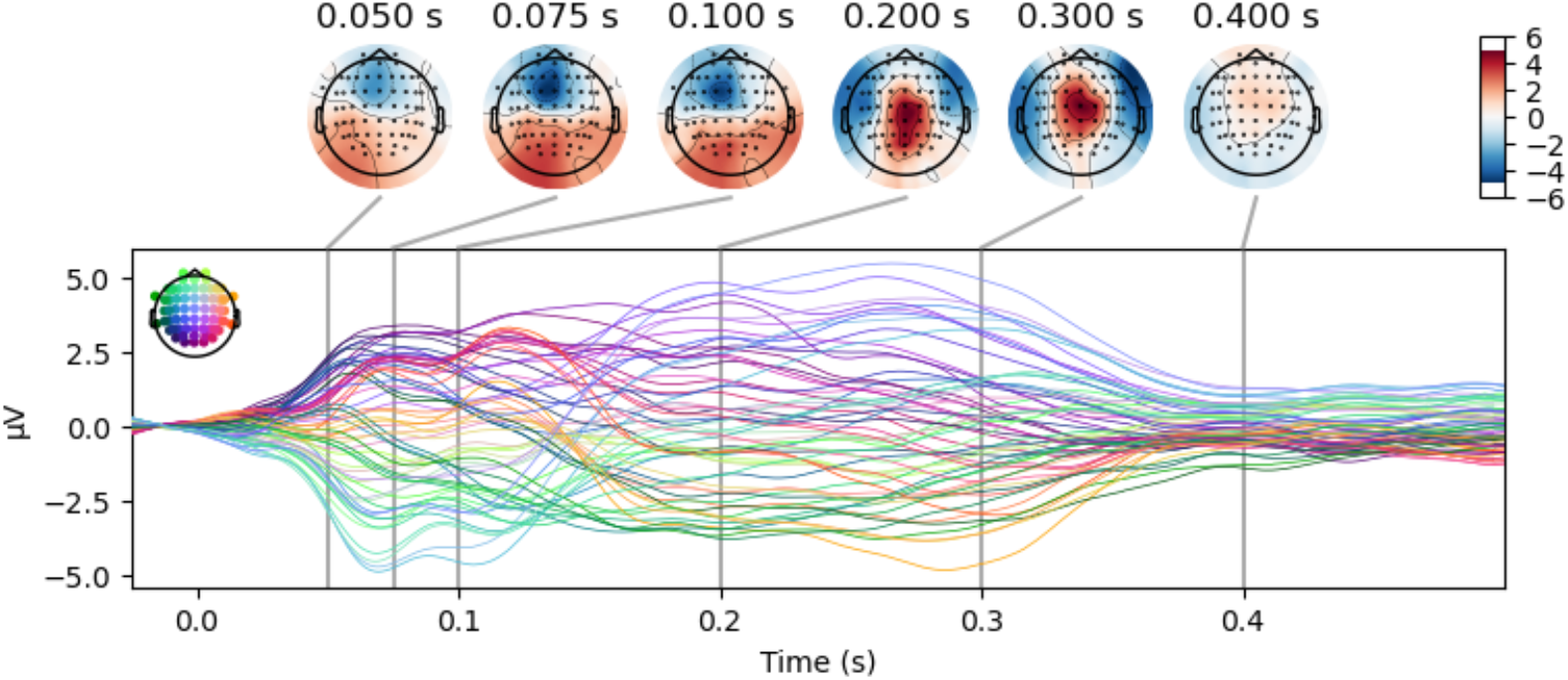
The grand-average evoked EEG response to muscle stimulation. The depicted waveform was computed by averaging the preprocessed (1-30 Hz filtered) data across stimulation trials within each subject, and then averaging the resulting subject-level EEG responses to obtain a group-level average.

**Figure 5:**
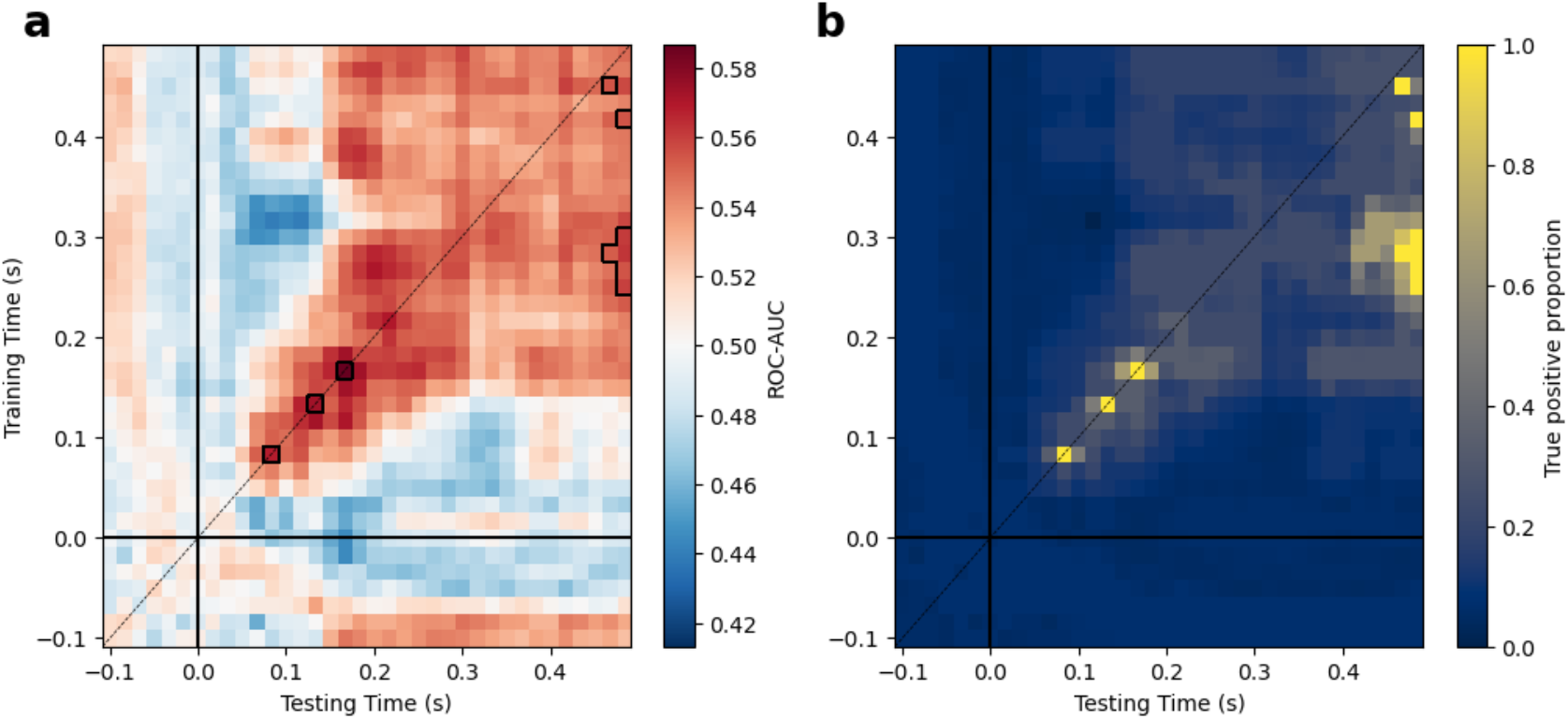
Temporal generalization of neural patterns predicting sense of agency. (a) Classification performance (ROC-AUC) for decoding subjects’ judgement of agency for individual muscle movements, cross-validated across subjects and across time. Results are shown for all (train-time, test-time) pairs to visualize the temporal dynamics of patterns that predict SoA. Above-chance decoding only near the diagonal reflects neural patterns which predict agency only transiently, whereas above chance decoding far off-diagonal reflects patterns which are sustained over time. Thus, patterns predicting agency appear to transition from transient to sustained dynamics around 170 milliseconds following stimulation. (b) Lower bounds on the true-positive proportion (TPP) within clusters, computed across all clustering thresholds. The value represented at each (train-time, test-time) pair is the highest TPP of any cluster in which that pair is included, thus larger values reflect greater certainty in the localization of effects.

**Figure 6:**
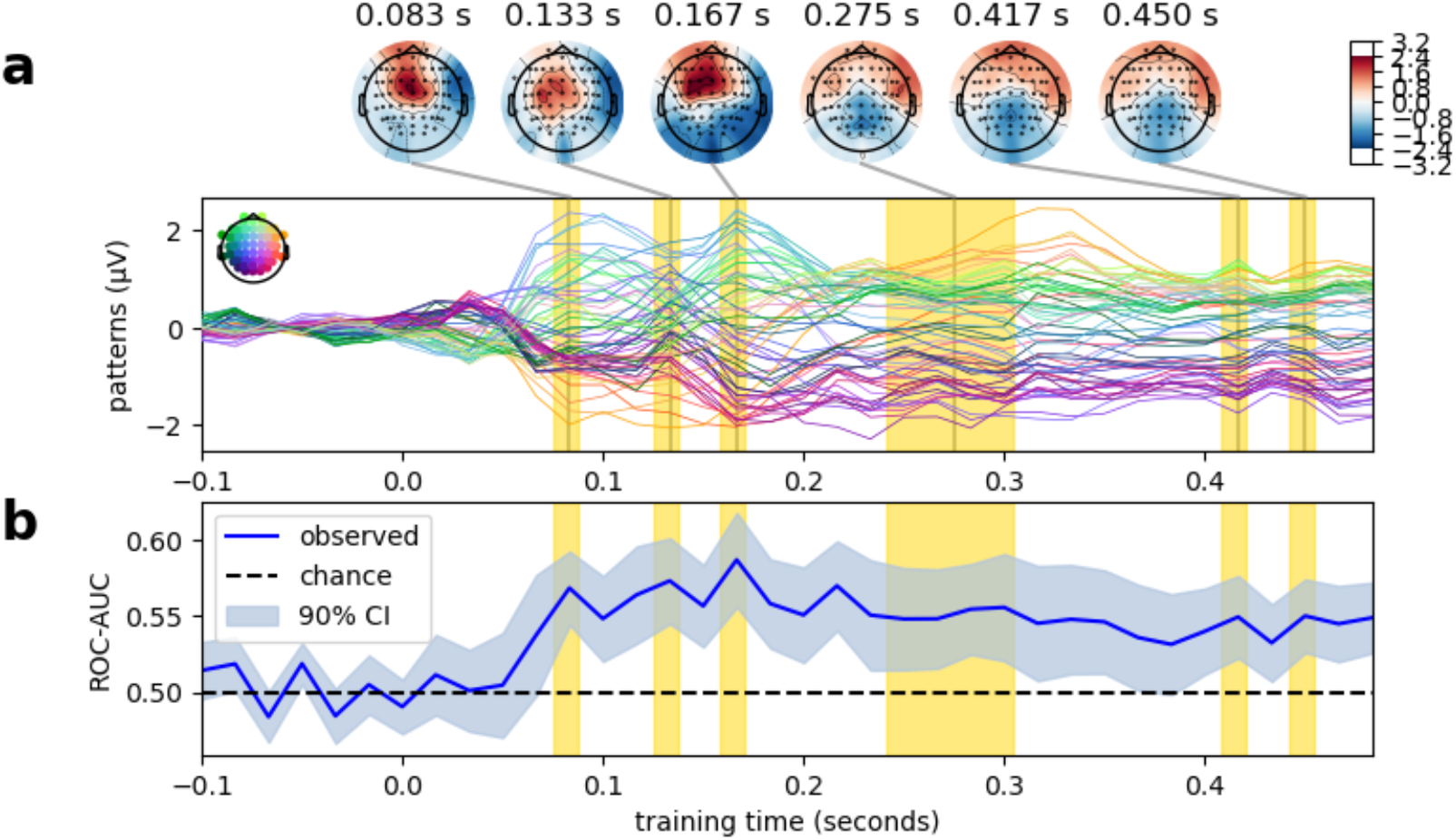
Voltage patterns that predict sense of agency. (a) The EEG topographies that the linear classifiers trained at each timepoint select for, reconstructed by inverting the trained classifier parameters using Haufe’s trick (Haufe et al., 2014). (b) The decoding performance when testing at each train time (identical to the values on the diagonal in Fig. 5a). Training times are highlighted in yellow if included in a cluster with TPP > 0.95 at any test time (Fig. 5b).

The earliest such cluster occurs 83 ms after the onset of muscle stimulation (adjusted threshold: 𝑝 < 4.5 × 10^45^). This is substantially earlier than previous studies have localized the earliest predictors of agency judgements (see *Discussion*), which may reflect a distinct role of low-level sensorimotor processes in agency judgments pertaining to the musculature itself, but less so to downstream sensory consequences of action. When comparing the patterns our decoding model selects for (see Figure 6) to the average evoked response (see Figure 4), one notes that the polarity of the pattern that predicts SoA is opposite the average response, indicating that the classifier would predict a self-agency judgment as the result when the sensory response is suppressed—a finding consistent with sensory attenuation (Voss et al., 2006). Classifiers trained at earlier times do not generalize to predict SoA at later times (see Figure 5), indicating that early prediction likely reflects a sequential chain of transient representations during sensorimotor processing (King and Dehaene, 2014). Later in the epoch, however, the temporal dynamics of the predictive patterns change to reflect a single, sustained neural signature that predicts SoA starting by at least 250 ms after stimulation and persisting at least until the end of the epoch (𝑝 < 0.003).

### 3.3. Fractal complexity of brain activity predicts agency judgements

Notably, trial-by-trial fractal dimension predicted SoA at almost every electrode (see Figure 7), reaching an ROC-AUC of 0.614 at electrode C1 (adjusted threshold: p < 0.027), even after the current source density transformation of the EEG signal was applied to attenuate the effects of volume conduction (see Methods). This suggest that the (local) complexity of the brain activity is increased uniformly throughout cortex following muscle movement when that movement is perceived as self-caused (as compared to when it is not perceived as self-caused). This is consistent with the previous observation that neural activity corresponding to self-generated percepts has a higher fractal dimensio**n** (Ibáñez-Molina and Iglesias-Parro, 2014).

**Figure 7:**
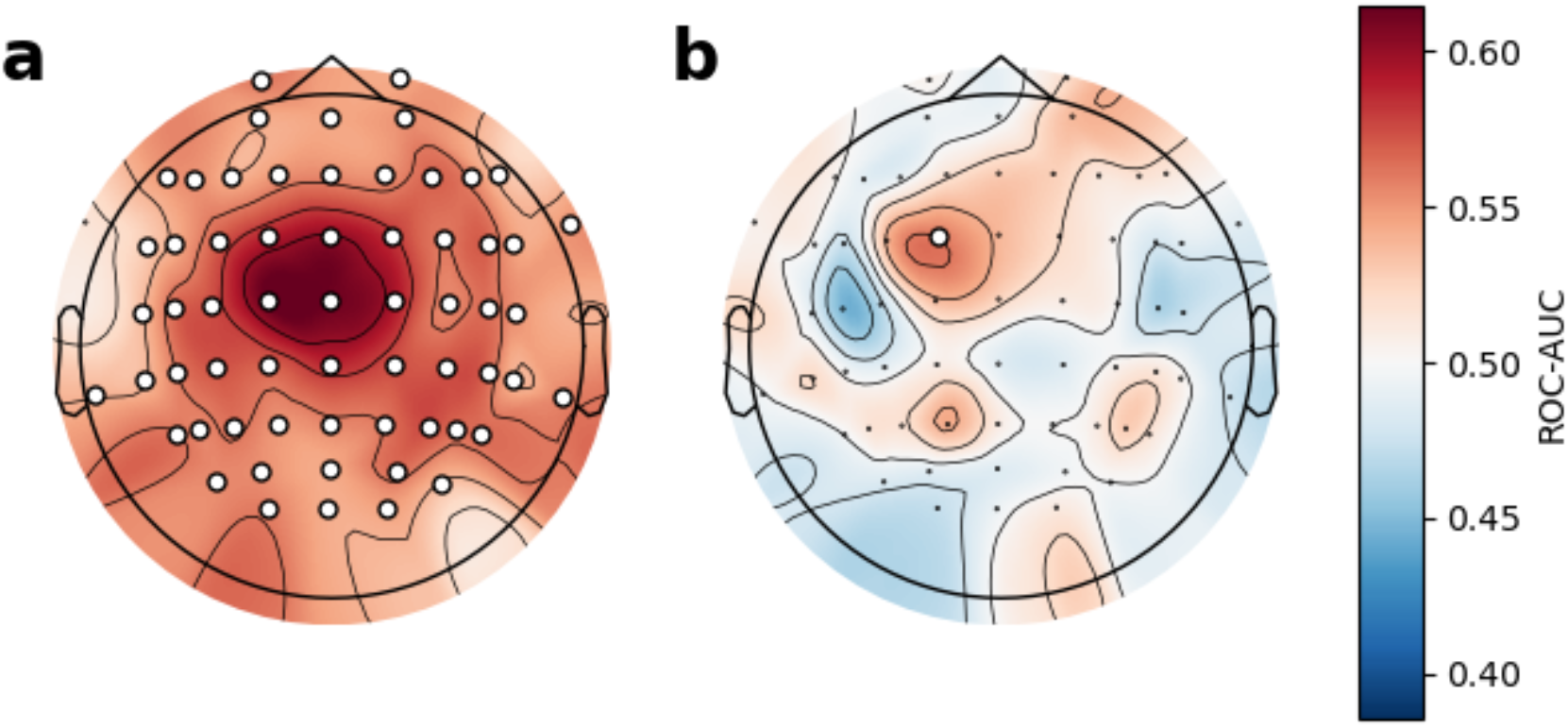
Classification performance for predicting trial-by-trial sense of agency from single-electrode fractal metrics. (a) Classification performance for Higuchi fractal dimension. (b) Classification performance for the Hurst exponent. Electrodes included in clusters in which the true positive proportion exceeds 95% are marked with white.

On the other hand, the Hurst exponent only predicted SoA at a single electrode at position FC1 (ROC-AUC = 0.559, p = 0.0006), located above cortical regions involved in motor control and planning, contralateral to the arm in which stimulation occurred (though we did not vary the arm used for stimulation, so we would caution against interpreting this as a strictly contralateral effect, though it is suggestive). This finding suggests a much more selective modulation of long-range temporal dependencies, such that the activity of specific frontocentral cortical regions becomes globally less to local perturbations in the absence of SoA—or, conversely, frontocentral areas are more sensitive to inputs in the presence of SoA. Notably, since the Hurst exponent (albeit only in one electrode) and fractal dimension both positively covary with agency, defying the strictly inverse relationship they would show in a strictly self-similar time series (see *Methods 2.4.3*), the EEG response to muscle movement appears to depart from (full) scale-freeness, at least over FC1. This divergence would allow the local complexity of and the temporal persistence of perturbations to neural activity to be modulated independently (see *Discussion*).

## 4. Discussion

Our findings advance our understanding of how the sense of agency (SoA) is generated in the brain, with important implications for the relationship between conscious self-awareness and unconscious self-referential processing. In particular, the time course neural activity predicting SoA in response to muscle stimulation is more consistent with classical sensorimotor monitoring accounts (Wolpert et al., 1995; Blakemore et al., 2000) than previous studies have shown comparing the neural responses to self- and other-caused tones (Kühn et al., 2011; Timm et al., 2016). While results still leave room for common downstream correlates of agency, they suggest that early responses differentiating self and other may be more *modality specific* than previously thought.

In the comparator model of SoA, originally imported from the motor control literature (Wolpert et al., 1995), sensations are compared to the intended or predicted sensory consequences of actions, and then congruent feedback is deemed self-caused and incongruent feedback externally caused (Feinberg, 1978; Frith, 1987; Gallagher, 2000). Since it is well documented that early sensory responses, especially those that are predictable, are attenuated during movement (Blakemore et al., 2000), it seemed plausible that the same machinery could parsimoniously account for conscious self-other discrimination. While this simple model is still the basis of most modern accounts of SoA, it is now understood that the mechanisms of conscious SoA diverge from low-level sensorimotor monitoring (Synofzik et al., 2008; Frith, 2012; Zaadnoordijk et al., 2019; Press et al., 2023).

To this effect, recent studies using typical paradigms, which probe the perception of causality between a button press and subsequent tone (i.e., “broad” sense of agency over action outcomes), have failed to find a relationship between the neural processes which would be affected by low-level sensorimotor monitoring – that is, the early, pre-conscious (< 200 ms) response to sensory stimulation – and conscious SoA (Voss et al., 2006; Kühn et al., 2011; Ohata et al., 2020). Timm and colleagues report a full dissociation, showing that comparator-model-like suppression of early responses to self-caused sensation occur in both the presence and absence of SoA (Timm et al., 2016). Since decades of research tell us early (< 200 ms) sensory responses reflect preconscious, rather than conscious, processing of the sensory stimulus (Libet et al., 1967; Sergent et al., 2005; Dehaene and Changeux, 2011), these findings have been interpreted as meaning that temporally early “exogeneous” neural responses (i.e., those that are a direct consequence sensory input) do not inform agency judgements, but later “endogenous” neural responses (e.g., P3 component) associated with conscious attention do (Kühn et al., 2011). None of these authors argue against the general idea of a comparator, but rather suggest that the comparison takes place at a higher level of abstraction than in the low-level sensorimotor monitoring used to guide motor learning (Wolpert et al., 2011).

In contrast, we find patterns in the early sensorineural response to stimulation predicts SoA even when that sensation was not actually self-caused, as we exclusively analyzed trials in which movements were caused by EMS. The critical difference is that we measured the neural response to muscle stimulation, and subjects made agency judgements about *the muscle movement itself* rather than a downstream consequence of movement. Thus, the mechanisms that give rise to narrow SoA over the musculature may overlap with basic sensorimotor processing more than those mechanisms that give rise to SoA over action outcomes more far removed from a subject’s motor intention (Charalampaki et al., 2022). Previous work manipulating bodily agency by altering the visual feedback from movement (leaving somatic feedback channels intact) has primarily used fMRI (Tsakiris et al., 2010; Abdulkarim et al., 2023) or EEG methods lacking the temporal resolution of the present approach (Kang et al., 2015); consequently, it is not totally clear whether our very early (preconscious) decoding results differ from previous findings merely because of our focus on SoA over body movements or because we additionally perturbed somatic (not just visual) feedback channels. Regardless, our data support the view that the earliest (pre-conscious) correlates of conscious SoA may differ based on context (i.e., what is one being asked to make a judgement about?), modality (e.g., proprioceptive or auditory), or level of abstraction.

However, it is worth noting that the earliest neural correlates of agency are not the end of the story. Indeed, the comparator model for SoA has largely been usurped by dual-process models in which the outcome of an initial comparator process is integrated with prospective, prior information to produce a final agency judgement (Synofzik et al., 2008; Haggard, 2017; Legaspi and Toyoizumi, 2019), and there is no clear theoretical for why or how multiple comparator processes taking place at multiple levels of abstraction may not be integrated into a single agency judgement. In fact, the shift we observe from transient to sustained patterns of neural activity predicting agency is quite consistent with that predicted by dual-process models of action processing (Del Cul et al., 2009; Charles et al., 2014). Specifically, the sustained nature of the predictive voltage patterns is consistent with a previously observed signature of high-level novelty/error detection that has been argued to require conscious awareness (Dehaene and King, 2016) and previously proposed to inform agency judgements (Kühn et al., 2011). An intriguing possibility then, which hybridizes the competing views proposed in the introduction, is that pre-conscious (roughly < 200 ms) predictors of SoA judgements will be context specific, but post-consciousness “neural correlates of self-awareness” integrate across modality-specific comparators. We do not manipulate awareness of action and outcomes here, so it is up to future work to test this hypothesis directly. Such investigations, which can compare sense of agency over actions with SoA over those actions’ downstream outcomes, are made possible by extending the paradigm we introduce here.

Further, both the fractal dimension – a measure of local signal complexity or “roughness” – and the Hurst exponent – a global measure of long-range correlation in a signal, indicative of how long a perturbation (e.g. sensory input) in the measured system would persist in time – were able to classify trial-by-trial SoA. However, the Hurst exponent was only predictive of SoA in a single frontocentral electrode, whereas fractal dimension was robustly predictive across the whole scalp. Both of these measures are often interpreted as reflecting a self-similarity or scale-free property of a time series, often appealing to theories of self-organized criticality as an explanatory framework (Churchill et al., 2016; Kardan et al., 2020; Zhuang et al., 2022). Indeed, the self-similarity interpretation has been invoked in explaining why the fractal dimension of neural activity corresponding to self-generated percepts is higher than that to external stimuli (Ibáñez-Molina and Iglesias-Parro, 2014). In a truly self-similar time series, however, fractal dimension and the Hurst exponent are strictly inversely related (Gneiting and Schlather, 2004); in contrast, both values positively covaried with SoA in the electrode in which we find Hurst was predictive. This finding suggests the neural response to muscle movement (as reflected in EEG) is not strictly self-similar, and so its complexity and sensitivity to perturbation can vary independently. While admittedly quite speculative, this observation may be interpreted as having functional importance, allowing sensorimotor cortical regions (which could possibly account for the frontocentral Hurst effect) to selectively modulate sensitivity to input, while overall cortex shows higher signal complexity with sense of agency.

In summary, while SoA has become a topic of increased attention in recent decades, most research in the area has focused on the experience of agency over downstream consequences of one’s actions as they affect the external world rather than the more basal experience of directing one’s own muscles (Haggard, 2008, 2017). We introduce the use of human-in-the-loop Bayesian optimization, in combination with electrical muscle stimulation, to experimentally manipulate the subjective experience of controlling the musculature. As we showcase here, this approach enables novel behavioral and neuroimaging investigations into the substrate of embodied self-awareness. Our results provide confirmatory evidence for the predictive relationship between low-level sensorimotor processes and SoA for muscle movements, which seems not to hold for the sensory response to action consequences (Dewey and Knoblich, 2014; Timm et al., 2016). While our findings suggest that early neural correlates of SoA may differ by context and modality, the transition from transient to sustained neural patterns that predict SoA in our data suggest at least two distinct neural processes contributing to agency judgments, as posited by dual-process theories of action selection and monitoring (Del Cul et al., 2009). This leaves open the possibility that modality-specific, pre-conscious predictors of SoA are still integrated into a single agency judgment downstream. Such a possibility could explain how information from multiple scales of biological organization are integrated into a unified experience of self, even if the mechanism of self-other differentiation differs across scales. We suggest that this hypothesis is a fruitful avenue of research for the emerging science of self-awareness.

## Author Contributions (CRediT Taxonomy)

**J.P.V.**: Conceptualization, Data curation, Formal analysis, Funding acquisition, Investigation, Methodology, Project administration, Software, Validation, Visualization, Writing - original draft, and Writing - review & editing. **P.L.**: Conceptualization, Funding acquisition, Methodology, Resources, Software, and Writing - review & editing. **H.C.N.**: Conceptualization, Funding acquisition, Resources, Supervision, and Writing - review & editing.

## Acknowledgements

P.L. and H.C.N. were supported by NSF NCS 2024923, and J.P.V. was supported by NSF GRFP DGE 1746045 and a Neubauer Family Distinguished Doctoral Fellowship.

